# Modeling two decades of elevational change in endemic bird densities in Hawaiʻi’s Kaʻū Rainforest

**DOI:** 10.64898/2026.08.20.746003

**Authors:** Philip T. Patton, Seth Judge, J. Andrew Royle, T. Scott Sillett

## Abstract

Protected areas are vital to the recovery of endangered species. Of the 24 remaining endemic passerines in Hawaiʻi, 16 species are endangered or critically endangered. Yet protected areas in the archipelago are changing as a result of climate change and biological invasions. For example, the non-native southern house mosquito (*Culex quinquefasciatus*), the primary vector of avian malaria (*Plasmodium relictum*), has been encroaching upward as higher elevations warm. How the ranges of endemic birds have also shifted their range upward in the Kaʻū Rainforest, the largest native forest on the Island of Hawaiʻi, is not well understood. We used hierarchical distance sampling to characterize how population density has changed from 2002 to 2024 for eight endemic bird species in the Kaʻū Rainforest. For five species, the most parsimonious model included an interaction between year and a quadratic elevation effect. Species that were most common at lower elevations in 2002, e.g., ʻApapane (*Himatione sanguinea*), tended to be most common at mid-elevations by 2024. Species that were already most common at higher elevations, such as ʻIʻiwi (*Drepanis coccinea*) and Hawaiʻi ʻĀkepa (*Loxops coccineus*), tended to decline in density. For example, ʻIʻiwi density at 1,530 m declined from 3.31 birds per ha (95% confidence interval [CI]: 2.67-4.11) to 1.34 birds per ha (95% CI: 1.05-1.71), and Hawaiʻi ʻĀkepa were completely extirpated from elevations below 1,530 m by 2024. Our results demonstrate how the elevational ranges of endemic species have shifted in response to climate change, biological invasions, and habitat degradation. Translating these results into conservation measures may require a more thorough investigation of the causal mechanisms of these range shifts with finer scale habitat data, under the same hierarchical modeling framework.

**LAY SUMMARY:**

- Protected areas such as National Parks, private lands managed for conservation, and state managed reserves have been essential to conserving Hawaiʻi’s native forest birds.
- Climate change and the cold-intolerant, invasive southern house mosquito, the primary vector of avian malaria, both threaten protected areas in Hawaiʻi. As the climate warms, avian malaria has been encroaching upwards to higher elevations.
- We found evidence that five of eight native forest bird species were shifting upward in elevation, perhaps in response to these dual threats, in Kaʻū Rainforest, the largest native forest in the Hawaiian Islands. The endangered Hawaiʻi ʻĀkepa has been extirpated from elevations below 1530m. Further, the vulnerable ‘I‘iwi has been extirpated from elevations below 1200m, and their density has declined by half at 1530m.
- Although more research could enhance understanding the mechanisms of this shift, our study supports wildlife managers in decisions related reducing the impact of mosquitos and climate change on native forest bird species within Hawaiʻi’s protected areas.

## INTRODUCTION

Area-based conservation, including protected areas and other effective area-based conservation measures, is widely regarded as one of the most effective strategies for recovering endangered species and conserving biodiversity. Recognizing this value, the Conference of the Parties to the Convention on Biological Diversity committed its member nations to conserving at least 30% of terrestrial and aquatic ecosystems by 2030 (Maxwell et al. 2020). Protected areas contribute to species recovery by maintaining habitat and simultaneously mitigating many threats recognized in the International Union for the Conservation of Nature (IUCN) Threat Classification Scheme, including residential and commercial development, agriculture, resource extraction, transportation infrastructure, biological resource use, and natural system modifications (Salafsky et al. 2025). Consequently, the effectiveness of area-based conservation has traditionally been measured by its ability to reduce these threats while maintaining the ecological conditions necessary for long-term species persistence (Watson et al. 2014). As environmental conditions and the relative importance of threats change over time, long-term ecological monitoring can provide information to help determine whether protected areas continue to meet the objectives for guiding adaptive management.

Few places illustrate the evolution of area-based conservation more clearly than the Hawaiian Islands. Hawaiʻi harbors one of the world’s most distinctive avifaunas, yet its endemic passerines are among the most threatened groups of birds. Since European colonization in the late 19^th^ century, 28 of the 58 described endemic species have gone extinct, and 21 of the remaining species are currently classified as threatened with extinction (IUCN 2025). Historically, widespread habitat conversion associated with agriculture, logging, and urban development motivated the establishment and expansion of protected areas that now encompass much of the remaining native forest across the archipelago (Scott et al. 1986; Banko and Banko 2009). Within these protected landscapes, state, federal, and private agencies prioritized fencing, feral ungulate removal, invasive plant control, and predator management (Hess and Jacobi, 2011; Hart et al. 2020). These investments protected and restored large tracts of native koa (*Acacia koa*) and ʻōhiʻa lehua (hereafter ʻōhiʻa; *Metrosideros polymorpha*) forest, improving habitat quality and facilitating the persistence of endemic forest bird populations across the islands (Hess and Jacobi, 2011; Hart et al. 2020).

Although protected areas continue to preserve native habitat, the primary drivers of decline for Hawaiian forest birds have shifted from habitat loss and degradation to invasive species and climate change operating within otherwise intact forests. The southern house mosquito (*Culex quinquefasciatus*), the primary vector of avian malaria (*Plasmodium relictum*) and avian pox (*Avipox* spp.), has become one of the greatest threats to endemic forest birds (van Riper et al. 1986). Following the introduction of mosquito-borne disease, endemic birds were largely extirpated from forests below 1,200-m elevation (Scott et al. 1986; Gorresen et al. 2009). Historically, cooler temperatures restricted mosquitoes to lower elevations (Atkinson et al. 2014), but climate warming has expanded suitable mosquito habitat upslope, eroding the high-elevation refugia that once protected many endemic species. Consequently, effective area-based conservation in Hawaiʻi increasingly depends on actively managing the ecological processes that drive extinction risk, including mosquito-borne disease transmission, depredation by non-native predators such as rats, cats, and mongooses, and the spread of Rapid ʻŌhiʻa Death, caused by the fungal pathogens *Ceratocystis lukuohia* and *Ceratocystis huliohia*, that threatens the native forest ecosystems where endemic birds persist (Barnes et al. 2018). Conservation success therefore depends on both protecting native forests and maintaining ecological conditions that sustain endemic bird populations.

The Island of Hawaiʻi provides a model system for evaluating how a changing climate is reshaping the distributions of endemic forest birds. As the youngest and largest island in the Hawaiian archipelago, it encompasses one of the broadest elevational gradients occupied by endemic passerines, extending from lowland wet forests to high-elevation montane woodlands. Because temperature influences the distribution of mosquitoes and avian malaria, this gradient provides a natural laboratory for detecting climate-driven shifts in bird distributions over time. Within this landscape, the Kaʻū Rainforest is particularly well suited for long-term study. Encompassing 35,000 ha of predominantly native forest on the southeastern slopes of Mauna Loa, it is the largest contiguous native forest in Hawaiʻi (Judge et al. 2024). The forest extends from approximately 600 m in wet mesic forest to over 2,300 m in montane woodland, encompassing much of the remaining elevational range occupied by endemic forest birds on the island. The landscape is managed collaboratively by the Hawaiʻi Department of Land and Natural Resources, Hawaiʻi Volcanoes National Park, The Nature Conservancy, and Kamehameha Schools, whose investments in habitat protection, ungulate exclusion, invasive species management, and ecosystem restoration have created one of the state’s largest protected areas.

Although climate-driven expansion of avian malaria is expected to reshape the elevational distributions of endemic forest birds, species differ in their susceptibility to disease and other stressors. The four honeycreepers (Fringillidae) currently recognized by the IUCN as threatened with extinction include the vulnerable ʻIʻiwi (*Drepanis coccinea*), the endangered Hawaiʻi ʻĀkepa (*Loxops coccineus*), ʻAlawī (Hawaiʻi creeper; *Loxops mana*), and ʻAkiapōlāʻau (*Hemignathus wilsoni*). These species are highly susceptible to avian malaria and therefore expected to exhibit the strongest contractions toward higher elevations as mosquito habitat expands (Atkinson et al. 2001; McClure et al. 2020). In contrast, ʻApapane (*Himatione sanguinea*) and Hawaiʻi ʻAmakihi (*Chlorodrepanis virens*) exhibit greater tolerance to avian malaria (Yorinks and Atkinson 2000; Woodworth et al. 2009; Atkinson et al. 2013) and continue to occupy portions of the lower elevation forest where disease transmission is common (McClure 2020; Bak et al. 2025). The Hawaiʻi ʻElepaio (*Chasiempis sandwichensis*), a monarch flycatcher (Monarchidae), is tolerant to avian malaria, but especially vulnerable to avian pox and depredation by rats (VanderWerf et al. 2006, 2009). Similarly, ʻŌmaʻo (*Myadestes obscurus*), a thrush (Turdidae), is also tolerant to avian malaria (Atkinson et al. 2001); changes in its density and distribution may instead reflect habitat degradation, non-native predators, or other ecological processes. Consequently, comparing long-term elevational changes among multiple species provides an opportunity to evaluate how different threats are reshaping endemic bird communities across one of Hawaiʻi’s largest protected landscapes.

To evaluate how endemic forest birds have responded to more than two decades of changing climate and evolving conservation challenges, we modeled species-specific changes in density across the elevational gradient of the Kaʻū Rainforest from 2002 to 2024. Our analysis included the eight endemic passerines described above. We estimated density with hierarchical distance sampling methods from nine years of point-transect surveys in the Kaʻū Rainforest. Our objective was to determine if the most parsimonious model for each species included an interaction between time and elevation and, if so, how the predicted density changed at different elevations over time (Gaya and Chandler, 2025). We validated these inferences by evaluating predictions on a holdout set for each species with Lorenz curves and calibration plots, underutilized tools in hierarchical modeling applications in ecology. These analyses provide a foundation for (1) evaluating how one of the world’s largest protected native forests functions under a rapidly changing threat regime, and (2) establish a baseline for assessing outcomes of emerging conservation interventions (e.g., landscape-scale mosquito suppression, predator control, habitat restoration) for Hawaiʻi’s remaining endemic forest bird communities.

## METHODS

### Data collection

Several state agencies, federal agencies, and non-governmental organizations—including the Hawaiʻi Division of Forestry and Wildlife, the National Park Service, the U.S. Geological Survey, The Nature Conservancy, and the Three Mountain Alliance Watershed Partnership—have conducted point-transect distance sampling surveys (hereafter, distance sampling surveys) in the Kaʻū Rainforest (Judge et al., 2024). The first effort was the Hawaiian Forest Bird Survey in 1976 (Scott et al., 1986). Our analysis focused on surveys from 2002, 2004, 2005, 2008, 2010, 2016, 2018, 2019, and 2024. The area covered by each effort, and the transects visited, varied by year (Figure 1). In general, each effort involved 8-minute distance sampling surveys at individual stations along transects. Some of these transects were denoted as “legacy” transects established in 1976, but others were randomly placed in the landscape each year. Stations on legacy transects were spaced 200-m apart, whereas stations on random transects were positioned 150-m apart. During each survey, researchers recorded the horizontal distance to any bird that they detected.

**Figure 1:**
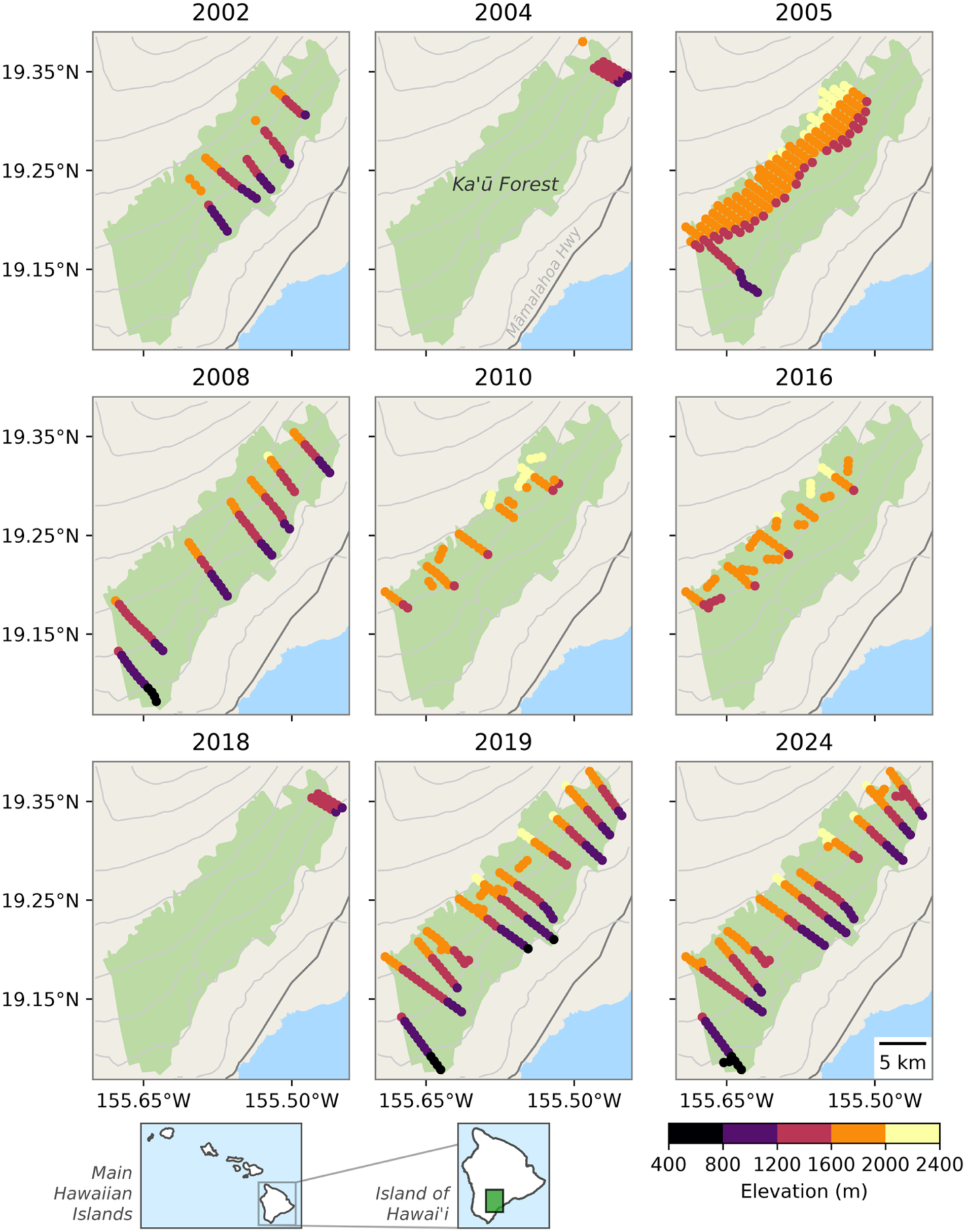
Locations and elevation of survey sites (colored dots) in Kaʻū Rainforest. To minimize overplotting, only sites that are at least 500 m apart are shown.

### Data preparation

We discarded any observations beyond 150 m from the observer and binned the remain set into distance intervals with a monotonic decline between intervals for all species (i.e., the number of birds detected in bin 1 was greater than that of bin 2, which was greater than that of bin 3, and so on). We then summed the total number of birds of each species within each distance bin at each “site”. We defined a “site” as a unique combination of year, survey replicate, transect, station number, and transect type (i.e., legacy or random) as an independent site. In other words, we treated temporal replications of surveys at a location as independent samples. Occasionally, stations on a transect were moved, but the “legacy” version of the station was still sampled. We treated these as independent sites.

We considered multiple covariates that might affect detection and density. Detection covariates were: year (categorically coded), canopy cover (categorical with levels: [very scattered, scattered, open, closed]), canopy height (ordinal with values: [0, 1, 2, 3]), wind-level (ordinal with values: [0, 1, 2, 3]; corresponding to Beaufort), time (minutes since midnight), a quadratic effect of time, wind-level, and percent cloud cover. Density model candidates were: *x* and *y* coordinates of survey station,, an interaction between *x* and *y* coordinates, linear and quadratic effects of site elevation, year (categorically coded, linear, and quadratic), an interaction between year and a quadratic effect of elevation (i.e., using R formula notation *year * elevation + year * I(elevation^2)*) in R software v. (R Core Team, 2026), canopy cover (categorical with levels: [very scattered, scattered, open, closed]), koa presence (categorical with levels: [present, absent]), and canopy height (ordinal with values: [0, 1, 2, 3]). We standardized continuous covariates (elevation, UTM coordinates, and survey start time) using z-score standardization. We imputed missing covariate values using the following approach: missing canopy cover, canopy height, and koa presence values were assigned from the nearest station with complete data, whereas missing weather variables (wind, cloud cover, rain, and gust) were assigned the modal value for that survey day.

### Hierarchical distance sampling

We estimated species-specific densities using hierarchical distance sampling models (Royle et al. 2004, Sillett et al. 2012) implemented in the R package unmarked (Fiske and Chandler, 2011, Kellner et al., 2023). To do so, we randomly split the data for each species such that 80% of the sites would be used for training, i.e., model fitting and selection, and 20% of the sites would be used for validation. As a result, the test and training sites were different for each species. We performed model selection in a two-stage approach. First, we performed forward model selection with the detection model, assuming constant density across sites. In the second stage, we performed forward model selection with the density model, assuming the best fitting detection model from the first stage. At each step of the forward selection, we fit models by adding each remaining candidate term and selected the term that yielded the lowest Akaike information criterion (AIC; Burnham and Anderson 2002) on the training data, provided the improvement exceeded ΔAIC > 2 relative to the current model. We excluded redundant terms from the candidate pool, e.g., a duplicative linear elevation effect when the quadratic effect had already been selected (note that the quadratic was always paired with a linear effect). We excluded models with singular Hessian matrices and models that produced estimates whose standard errors exceeded 5 on the link scale, suggesting poor identifiability. We assumed a half-normal detection function for all species.

### Model validation

We calibrated the model’s predictions by evaluating its fit on the holdout set with two graphical devices: calibration plots and Lorenz curves (Blöchlinger, 2012). The calibration plots helped us evaluate the model’s ability to capture the relationship between elevation and density over time. To build the plot, we predicted the number of birds detected in each distance bin for every site in the holdout set. We averaged (mean) the predictions and observations across distance bins for each site. Further, we averaged (mean) the predictions and observations by “elevation bin”, which corresponded to the nearest value on the standardized scale, which spanned [-2, -1, 0, 1, 2] corresponding to true elevation values (in meters) of [858, 1,200, 1,530, 1,870, 2,210]. This arrangement allowed us to directly calibrate our inferences about a certain elevation band and year for each species and to identify any weaknesses in our assessment of the relationship between elevation and density over time.

## RESULTS

Surveyors detected 36,904 birds of the eight endemic species at 2,697 sites from 2002 to 2024. Most of these detections were ʻApapane (22,378 total), followed by Hawaiʻi ʻAmakihi (6,523), ʻŌmaʻo (5,091), ʻIʻiwi (1,799), and Hawaiʻi ʻElepaio (470). The least detected species were the ʻAkiapōlāʻau (52), ʻAlawī (281), and Hawaiʻi ʻĀkepa (310). The smallest distance bin that would ensure a monotonic decline in counts across all species was 50 m, resulting in three bins: (0, 50], (50, 100], and (100, 150] (Figure 2). The training/validation split resulted in 2,186 sites for training and 511 sites for validation.

**Figure 2:**
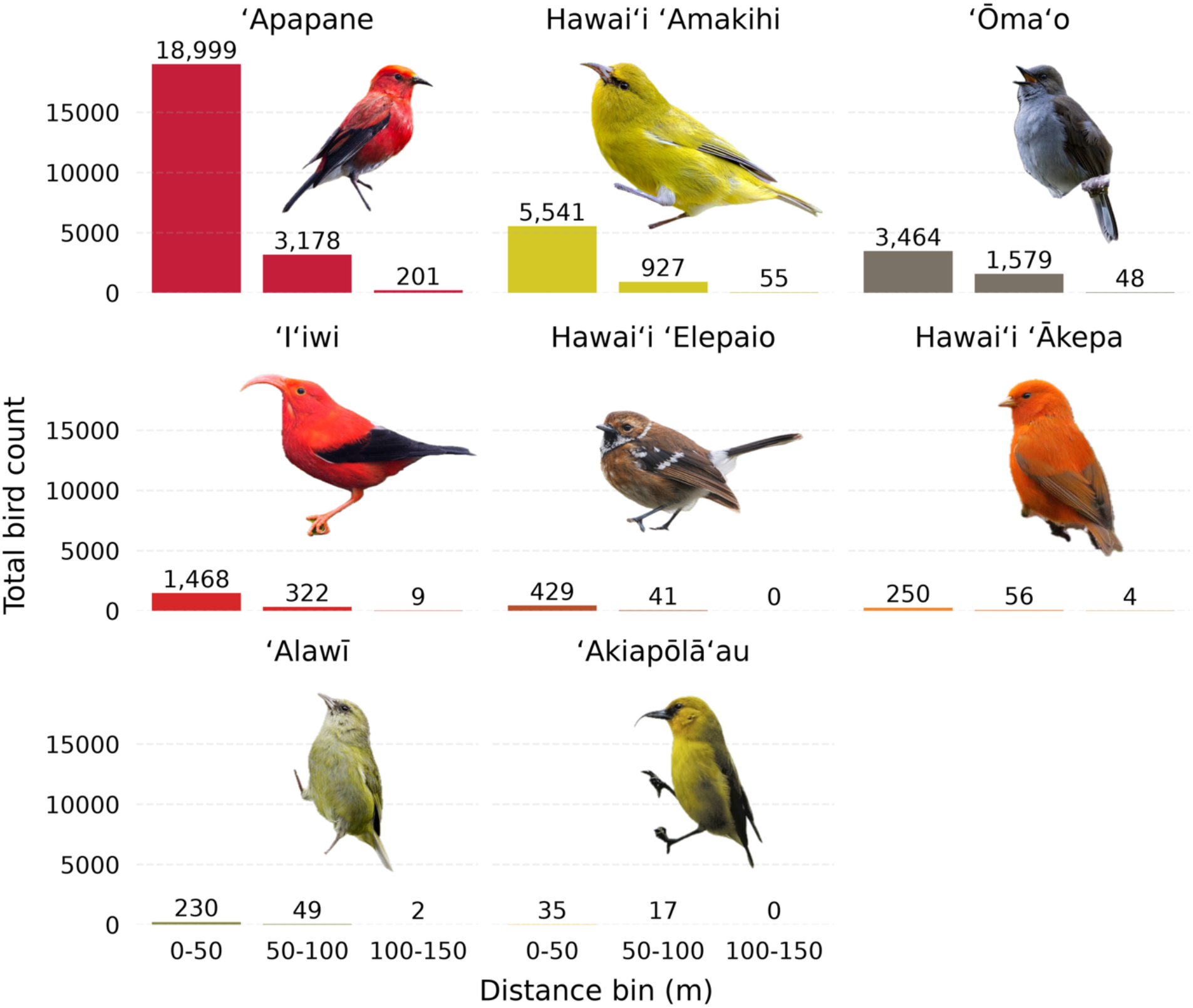
Total number of detections (colored bars) for each species within each distance bin.

The model selection procedure produced distinct models for each species, although the detection and density models included similar terms for many species (Tables 1 and S1). For example, the detection model for seven of eight species included an effect of canopy cover, and for six species, the detection model included the time of the survey. The density model for each species included an interaction term between the *x*-coordinate and the *y*-coordinate of the station. Additionally, the density model for seven species included some year effect. For Hawaiʻi ʻAmakihi, this was a categorically coded year effect. For Hawaiʻi ʻElepaio, this was a quadratic effect of year. For the other five species, the density model was an interaction between year and a quadratic effect of elevation. As a result, the best fitting model for these five species suggested that the effect of elevation on density was changing over time. For example, areas of highest ʻApapane density shifted from lower elevations (669 m) in 2002 to middle elevations (1,411 m) in 2024 (Figures 3 and 4). Note we omitted ʻAlawī, whose model of best fit included a very subtle elevation by year interaction, from Figures 3, 4 and 6 for clarity; this interaction may be a random artefact of model selection. For ʻIʻiwi, the areas of highest density in 2002 were already roughly one standard deviation above average elevation (1,715 m) and increased little by 2024 (1,816 m) (Figures 3 and 4). Note that predicted Iʻiwi density at those elevations declined from 4.90 birds per ha (95% CI: 4.07 - 5.89) to 3.67 birds per ha (2.79 - 4.81). Two lower elevation species ʻApapane and ʻŌmaʻo, appeared to shift upward, whereas two high elevation species, Hawaiʻi ʻĀkepa and ʻIʻiwi, moderately shifted upward (Figure 4).

**Table 1:** Terms included in the most parsimonious model for each species.

| Term | ‘Ōma‘o | ‘Apapane | Hawai‘i<br>‘Amakihi | ‘I‘iwi | Hawai‘i<br>‘Ākepa | Hawai‘i<br>‘Elepaio | ‘Alawī | ‘Akiapōlā‘au |
| --- | --- | --- | --- | --- | --- | --- | --- | --- |
| **detection** |  |  |  |  |  |  |  |  |
| factor(year) | ✓ | ✓ | ✓ | ✓ |  | ✓ |  |  |
| Canopy_Height | ✓ | ✓ | ✓ | ✓ |  | ✓ |  | ✓ |
| Gust | ✓ |  |  | ✓ |  |  | ✓ |  |
| Canopy_Cover | ✓ | ✓ | ✓ | ✓ | ✓ | ✓ | ✓ |  |
| time | ✓ | ✓ | ✓ | ✓ | ✓ |  | ✓ |  |
| Wind | ✓ |  | ✓ |  |  | ✓ | ✓ |  |
| **density** |  |  |  |  |  |  |  |  |
| elev * year | ✓ | ✓ |  | ✓ | ✓ |  | ✓ |  |
| l(elev^2) * year | ✓ | ✓ |  | ✓ | ✓ |  | ✓ |  |
| Canopy_Height | ✓ | ✓ |  | ✓ |  | ✓ |  |  |
| x * y | ✓ | ✓ | ✓ | ✓ | ✓ | ✓ | ✓ | ✓ |
| Canopy_Cover | ✓ | ✓ | ✓ | ✓ |  |  |  |  |
| factor(year) |  |  | ✓ |  |  |  |  |  |
| elev |  |  | ✓ |  |  | ✓ |  | ✓ |
| l(elev^2) |  |  | ✓ |  |  | ✓ |  | ✓ |
| year |  |  |  |  |  | ✓ |  |  |
| l(year^2) |  |  |  |  |  | ✓ |  |  |

**Figure 3:**
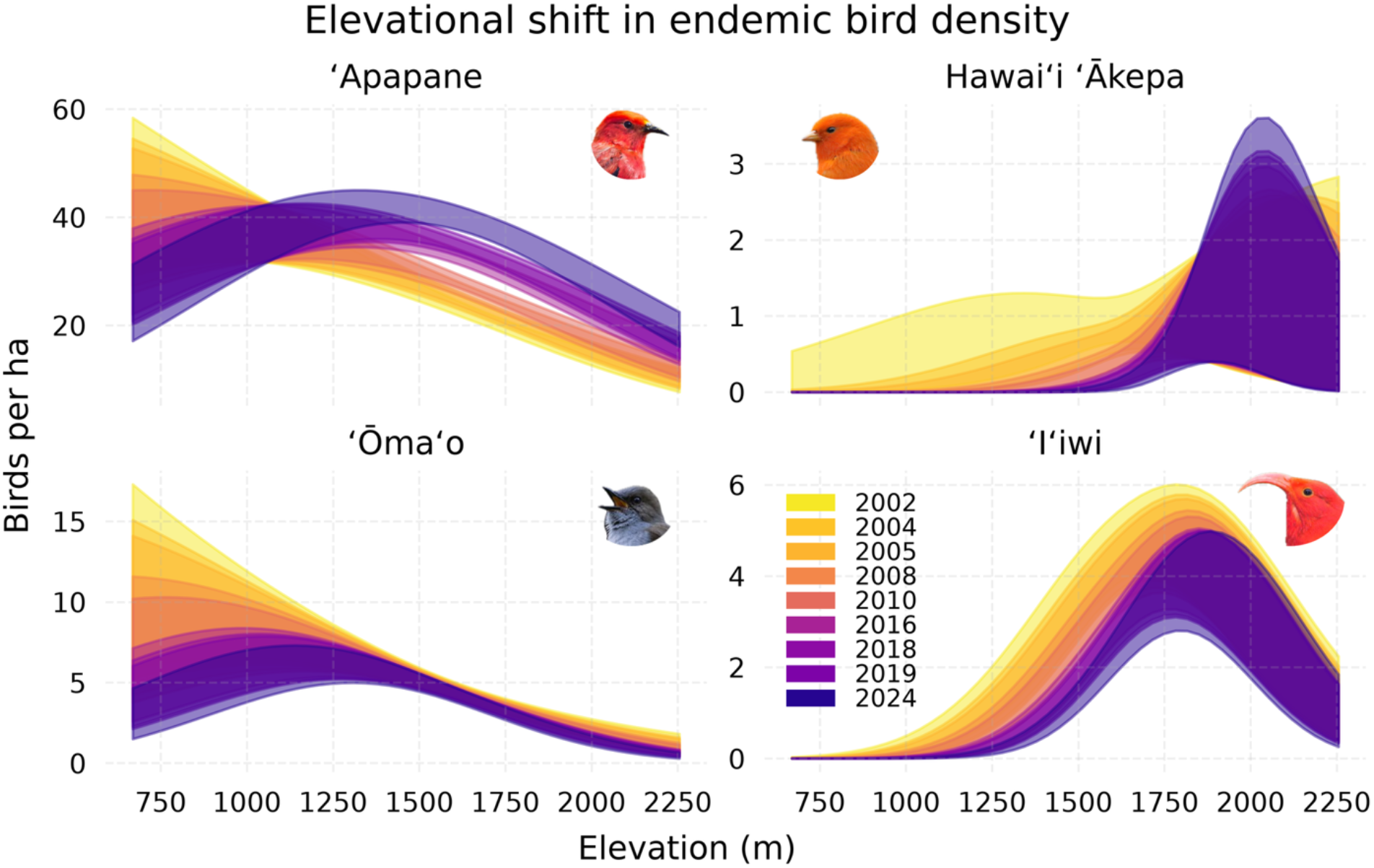
Changing effect of elevation on predicted density of four endemic species (colored ribbons) over time. Each ribbon represents the 95% confidence intervals, with darker colors represent later years. Predictions are conditional on average values of continuous covariates and modal values of categorical covariates.

**Figure 4:**
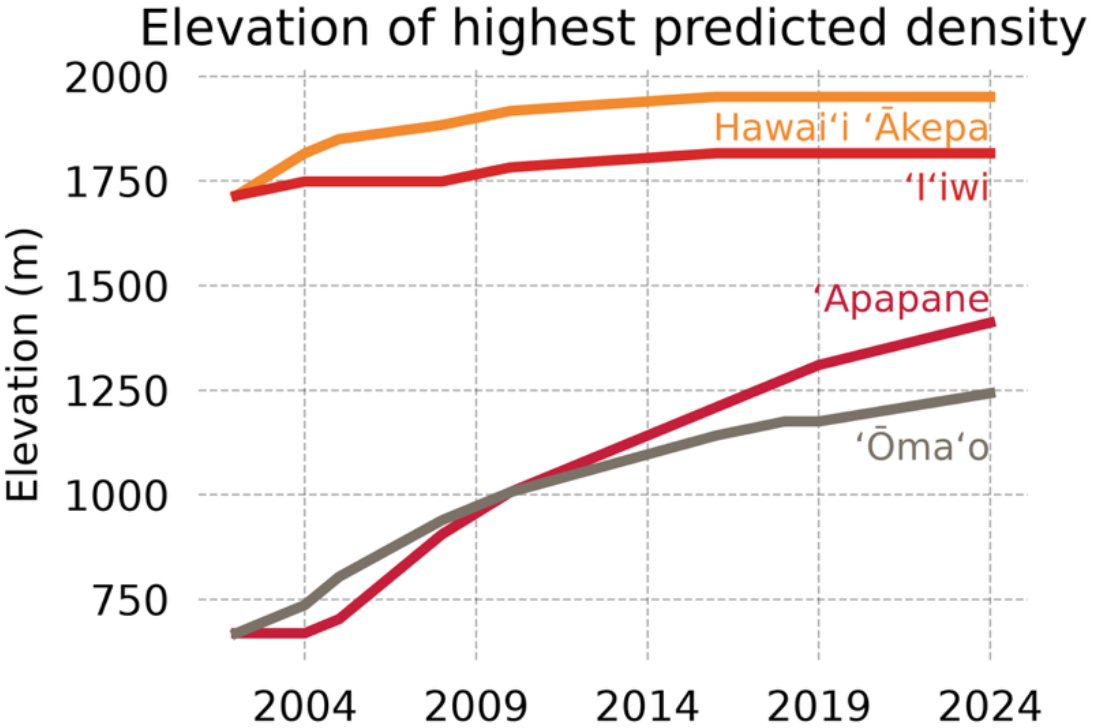
The elevation at which the density was predicted to be highest in each year (colored lines) for four species. For example, the density of ʻApapane was highest, all else being equal, at 669 m in 2002. In 2024, the density of ʻApapane was highest at 1411 m. Predictions are conditional on the average value of all continuous covariates, and the modal value of all categorical covariates.

**Figure 5:**
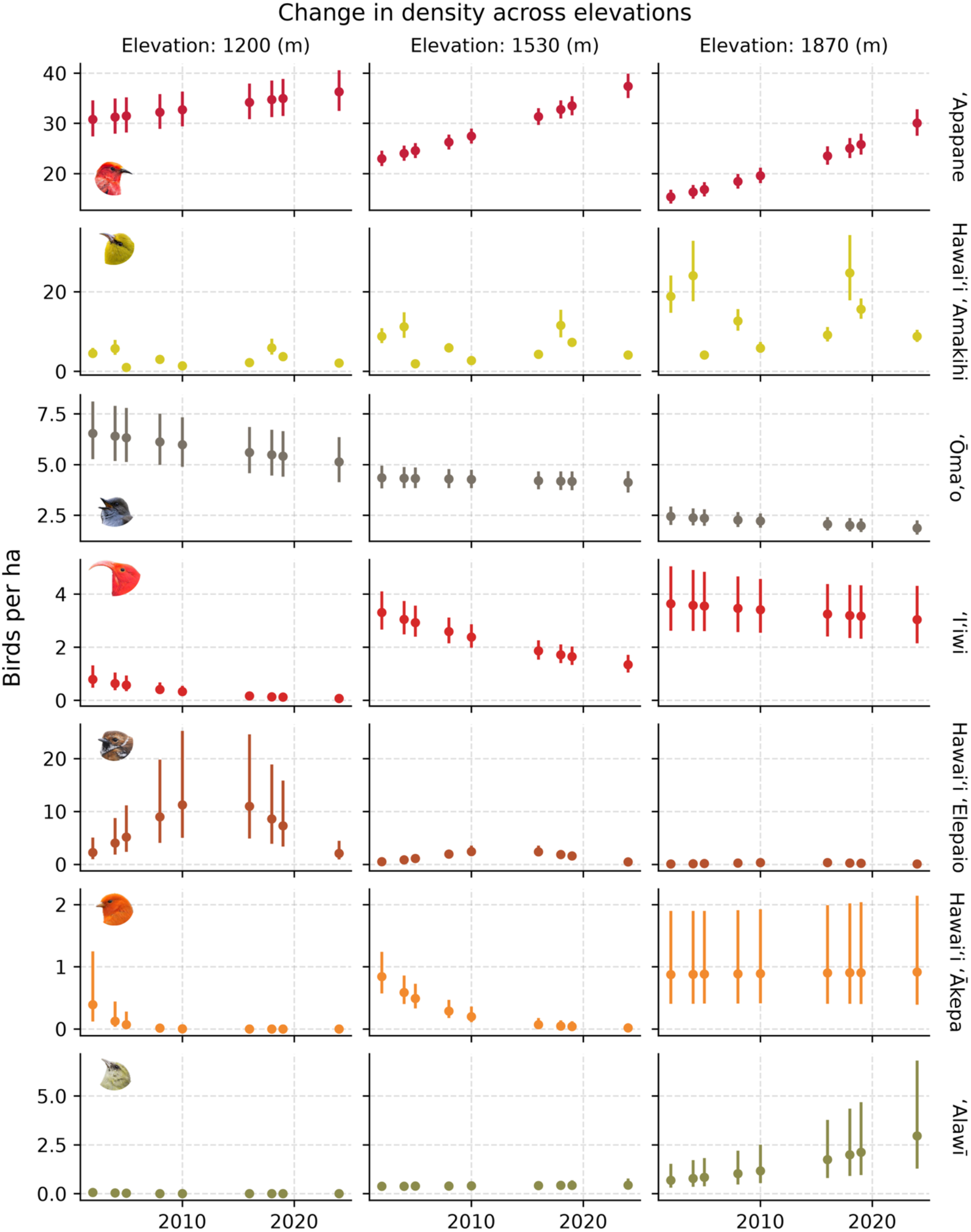
Predicted density at three elevations (birds per ha; colored dots) as a function of time, with 95% confidence intervals (colored lines). Elevations represent the mean elevation in the study area and plus or minus one standard deviation from the mean, Predictions are conditional on the mean value of other continuous covariates and the modal value of categorical covariates. Elevation values are the mean and one standard deviation above or below the mean.

**Figure 6:**
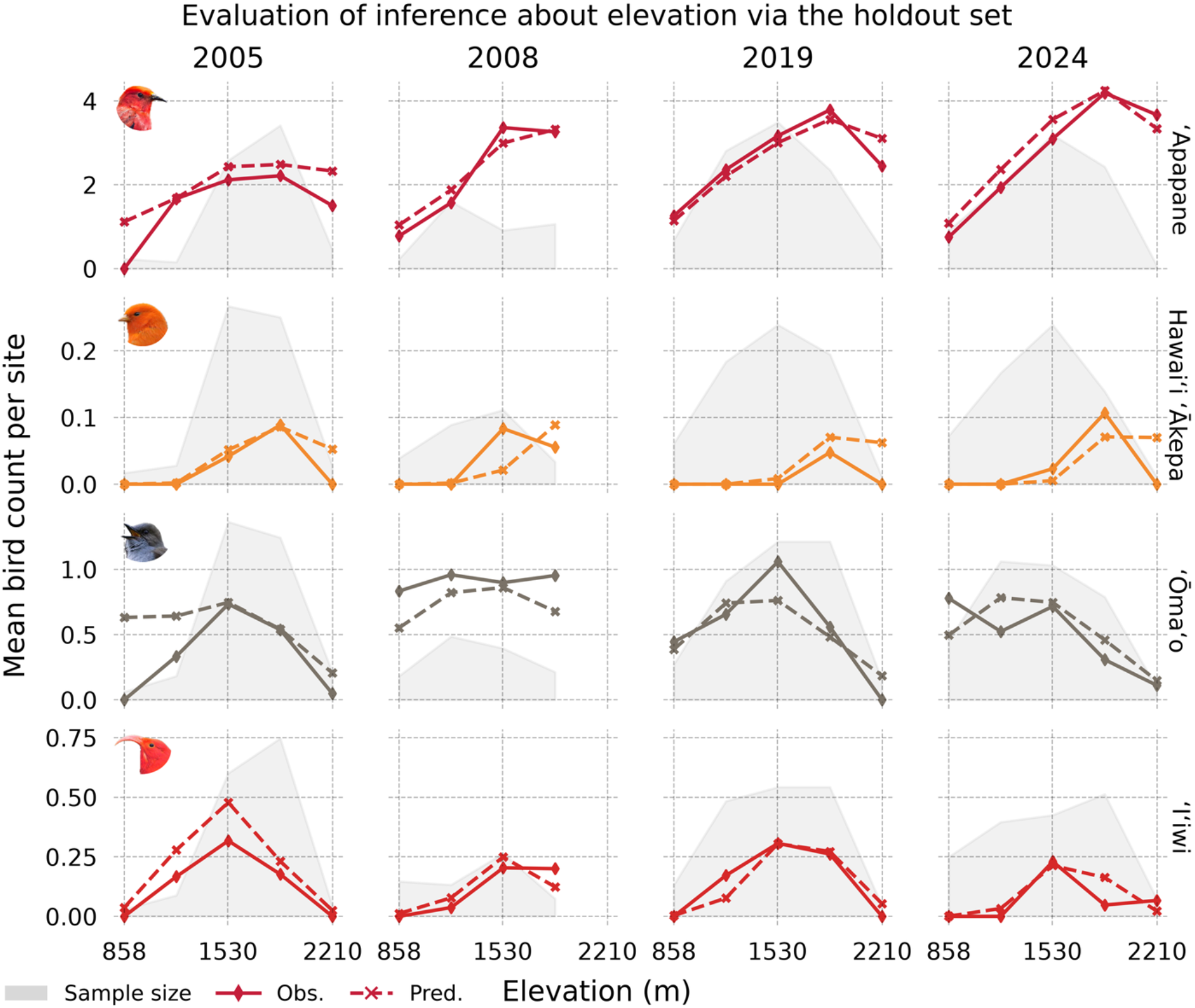
Average total observed (solid lines with diamonds) and predicted (dashed lines with x) number of birds detected at holdout sites. Holdout sites were allocated to one of eight equally spaced bins based on their elevation (akin to a histogram). Observed and predicted values were averaged for each bin. The number of sites in each bin is represented by the grey shaded area. Note that the test/train split was random for each species, hence, the slight variation in the sample size and x-axis by species.

**Figure 7:**
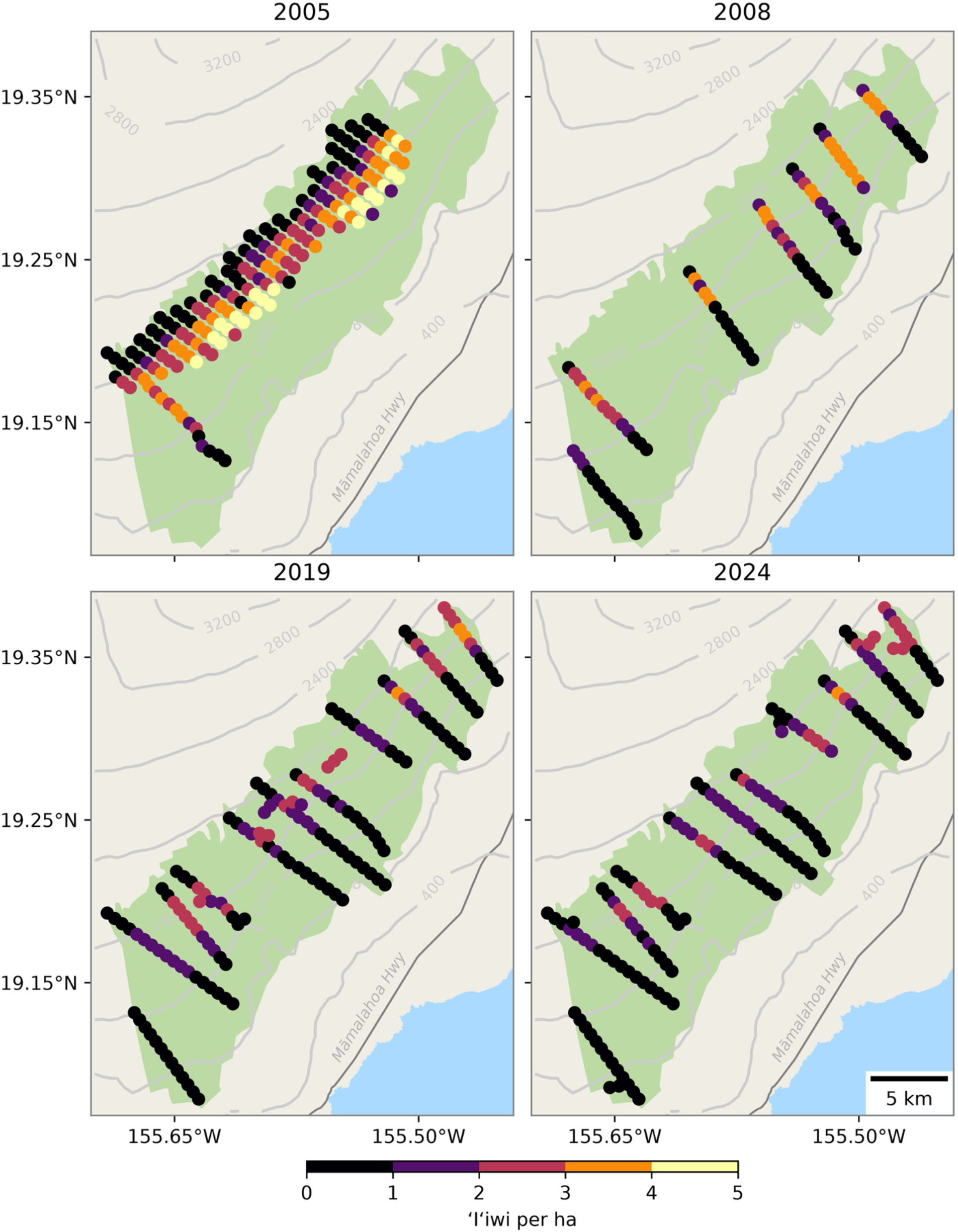
Predicted density (birds per ha) of ʻIʻiwi at survey points (colored points) from the four survey years with the most effort. Only points at least 500m apart are shown to minimize overplotting.

The change in density over time varied across species (Figure 5 and Table S2). Two species, Hawaiʻi ʻĀkepa and ʻIʻiwi, significantly declined in density from 2002 to 2024 at low and mid elevations (Figure 5) (95% confidence intervals for 2002 and 2024 did not overlap). For Hawaiʻi ʻĀkepa, the model predicted a decline from 0.39 birds per ha (0.12-1.25) to 0.00 birds per ha (0.00-0.00) from 2002 to 2024 at 1,200 m, and a decline from 0.84 birds per ha (0.57-1.24) to 0.01 birds per ha (0.00-0.07) at 1,530 m. Note that the mean elevation was 1,530 m and 1,200 m was one standard deviation below the mean. ʻIʻiwi declined from 0.79 birds per ha (0.48-1.32) to 0.07 birds per ha (0.04-0.14) at 1,200 m, and from 3.31 birds per ha (2.67-4.11) to 1.34 birds per ha (1.05-1.71) at 1,530 m. Neither species significantly declined at 1,870 m (95% confidence intervals for 2002 and 2024 overlapped). ʻŌmaʻo appeared to have declined at 858 m, i.e., two standard deviations below the mean elevation, from 8.27 birds per ha (5.56-12.3) to 3.59 birds per ha (2.28-5.43), but otherwise did not significantly change in density (i.e., overlapping confidence intervals). Hawaiʻi ʻElepaio, whose best model did not include an interaction between year and elevation, did not have a significantly different density in 2002 versus 2024 (95% intervals overlapped). However, it appeared to grow then decline in density at 1,200 m and 1,530 m (see below). Hawaiʻi ʻAmakihi significantly fluctuated in density between years, although without indicating overall growth or decline. The endangered ʻAlawī did not significantly grow or decline at 1,200 m or 1,530 m, yet appeared to grow in density at 1,870 m from 0.69 birds per ha (0.31- 1.53) in 2002 to 2.96 birds per ha (1.28-6.82) in 2024. Conversely, ʻApapane dramatically grew in density at 1,530 m from 23.0 birds per ha (21.5-24.6) to 37.4 birds per ha (35.0-40.0), and at 1,870 m from 15.3 birds per ha (14.0-16.8) to 30.1 birds per ha (27.6-32.8). Estimates of ʻAkiapōlāʻau density were small and highly uncertain (Table S2).

Predictive performance on the holdout set largely supported our inference about the effects of elevation and time on density (Figure 6). Predicted counts at different elevations over time closely matched observed counts for ʻApapane, supporting the inference about their decline at low elevations and dramatic growth at high elevations. For both Hawaiʻi ʻĀkepa and ʻIʻiwi, predictions generally matched observations with a few differences. For Hawaiʻi ʻĀkepa, the model tended to overpredict counts at elevations above average. The model generally predicted counts of ʻIʻiwi well except at mid elevations in 2005, where it overpredicted counts. Otherwise, predictions for both Hawaiʻi ʻĀkepa and ʻIʻiwi closely matched observations at elevations at or below 1,530 m, supporting inferences about declines at these elevations. For ʻŌmaʻo, the model overpredicted counts at below-average elevations before 2008, suggesting that the gentle decline shown in Figure 5 may be overstated. The model struggled to accurately predict counts of Hawaiʻi ʻElepaio across elevations in 2019 and 2024. The observed data offer a clue why: whereas 2008 (and 2005, to some extent) show a clear quadratic relationship between elevation and observed counts, the relationship is much more erratic in 2019 and 2024. This suggests that local factors beyond elevation may be influencing Hawaiʻi ʻElepaio counts.

## DISCUSSION

We estimated the density of eight Hawaiian endemic passerines across the Kaʻū Rainforest from 2002 to 2024. Six of the eight species were best described by models that included an interaction between year and a quadratic elevation effect, indicating that changes in density varied across the elevational gradient (Figures 3 and 4). The distributions of several species, including ʻApapane, shifted upward, with densities declining at lower elevations (∼850 m), and increasing at mid- (1,200-1,530 m) and high elevations (1,870-2,210 m; Figure 5; Table S2). Specialists such as ʻIʻiwi and Hawaiʻi ʻĀkepa continued to contract toward higher elevations, consistent with the upslope expansion of mosquito-borne disease (Samuel et al. 2015). Densities of ʻIʻiwi significantly declined at mid-elevations, whereas Hawaiʻi ʻĀkepa were extirpated from elevations below 1,530 m by 2024, but were stable at high elevations. We also observed significant declines of ʻŌmaʻo at lower elevations, despite this species being tolerant to avian malaria. We found evidence that the density of the endangered ʻAlawī increased at higher elevations over the study period. While the density at 1870m in 2024 was not significantly different the density at that elvation 2002, this does echo results from Hakalau National Wildlife Refuge in the northwest of Hawaiʻi Island (Hunt et al. 2026). Further, ʻAlawī remains restricted to elevations above 1,530 m. The endangered ʻAkiapōlāʻau remained nearly absent at low and mid-elevations and persisted at low densities (≤ 0.1 ± 0.6 SE birds ha⁻¹) in mature canopies codominated by ʻōhiʻa and koa. Overall, model predictions closely matched observed elevational patterns for several species, including ʻIʻiwi, Hawaiʻi ʻĀkepa, and ʻApapane (Figure 6). However, weaker predictive performance for Hawaiʻi ʻAmakihi and Hawaiʻi ʻElepaio suggests that stressors operating at finer spatial scales may exert a stronger influence on their distributions than elevation alone.

The strongest evidence for climate-driven expansion of avian malaria was observed with the mid-elevation declines of ʻIʻiwi. The species is highly susceptible to avian malaria, with a 90- 100% mortality rate for exposed birds (Atkinson et al., 1995). Historically, the upper limit of *Culex* mosquito distribution and malaria transmission was around 1,500-m elevation, where mean temperatures were approximately 14℃ (Atkinson et al., 2000; Ahumada et al., 2004). However, temperatures in Hawaiʻi have warmed significantly by 0.05℃ per decade over the last 100 years (McKenzie et al., 2019). Since 2002, densities of ʻIʻiwi at the 1,530-m elevation band declined by 60% to an estimate of 1.3 ± 0.17 SE birds ha⁻¹ in 2024. This species is nearly extirpated at the 1,200-m band, a pattern consistent across the main Hawaiian Islands (Paxton et al., 2013). In contrast, ʻApapane densities at 1,530 m increased by 60-65%, reaching a peak of 37.4 ± 1.25 SE birds ha⁻¹ in 2024, demonstrating a pronounced shift in the balance between the two nectarivores. The larger ʻIʻiwi has been observed aggressively defending and competitively excluding smaller honeycreepers such as ʻApapane (Carpenter, 1976). ʻApapane’s current dominance at mid-elevations suggests that it is exploiting foraging resources that ʻIʻiwi rely upon to sustain productivity.

Although the responses of ʻIʻiwi and the three endangered honeycreepers closely follow expectations for malaria-sensitive species, not all endemic birds exhibited patterns consistent with climate-driven disease alone. Several malaria-tolerant species also declined, particularly at lower elevations, suggesting that additional stressors are influencing populations within the Kaʻū Rainforest. These contrasting responses demonstrate that avian malaria, though the dominant threat for many endangered honeycreepers, cannot fully explain recent changes in Hawaiʻi’s forest bird community.

Fluctuations in densities of ʻŌmaʻo, Hawaiʻi ʻElepaio, Hawaiʻi ʻAmakihi, and portions of the ʻApapane population at low to mid-elevations are more consistent with sensitivity to habitat degradation and rat predation, rather than malaria-driven dynamics. Both the Hawaiʻi ʻElepaio and the endangered Oʻahu ʻElepaio (*C. ibidis*) have benefited from rat-control programs, where removal of black rats (*Rattus rattus*) has led to sharp increases in nest survival, fledging success, and the stabilization of subpopulations (VanderWerf et al., 2012; Banko et al., 2019). The Oʻahu ʻElepaio has responded positively to species-specific management, particularly in forests dominated by non-native avifauna, but we expect many native species to be vulnerable to rat predation and therefore to benefit from broad rat-control efforts in the Kaʻū Rainforest. For ʻŌmaʻo, we observed significant declines at low elevation bands, where density declined by 55- 60% from early years (2002-2005) to 3.6 ± 0.76 SE birds ha⁻¹ in 2024. This frugivore depends on the availability of native fruiting species (Wakelee and Fancy, 1999), yet invasion by weedy species, facilitated by ungulate disturbance, has degraded many low-elevation areas that remain unfenced in the Kaʻū Rainforest. In recent survey years, observers noted frequent pig signs at low elevations and occurrence of invasive plants such as *Psidium cattleianum*, *Passiflora tripartita* var. *mollissima*, *Cestrum nocturnum*, *Miconia crenata*, and others at the lowest stations, effectively restricting ʻŌmaʻo and other endemic birds to higher, less disturbed elevations. Furthermore, the wilt pathogen Rapid ʻŌhiʻa Death has spread into portions of the Kaʻū Rainforest (Vaughn et al., 2023) and is expected to negatively affect forest birds by causing habitat loss throughout their distribution (Camp et al., 2019).

### Statistical considerations

We also demonstrated an underutilized approach in hierarchical modeling in ecology, namely, evaluating predictions on a holdout set. Every year there are more long-term or large-scale ecological datasets, many of which were collected with hierarchical modeling in mind. Yet many applications still train models and validate predictions with the same dataset. This approach risks overfitting to the training data, especially with higher variance methods, such as generalized additive models (Miller et al., 2013). Moreover, it misses the opportunity to interrogate the model’s predictions regarding the scientific question of interest. For example, we were able to bolster our inference about changes in ʻIʻiwi and Hawaiʻi ʻĀkepa density, while highlighting areas of improvement for modeling Hawaiʻi ʻElepaio density (Figure 6). When coupled with thoughtful predictive modeling (Nichols & Cooch, 2025), the evaluating predictions on a holdout set can be a powerful tool for causal inference. One challenge for hierarchical models in ecology, where the processes of interest like density and abundance are unobserved, is designing performance metrics that disentangle observation from process. While challenging, researchers have devised clever fit statistics to tease apart latent and observed processes in hierarchical models in ecology (Glennie et al., 2019).

The density of Hawaiʻi ʻElepaio was not well-characterized by the covariates and relationships stipulated here. In fact, for all species, there were areas of improvement in terms of our ability to predict the number of birds counted at a station. This could be for challenges with modeling the detection process, or with modeling the density process. For example, we hypothesize that Hawaiʻi ʻElepaio density would be driven by understory structure, e.g., native plant composition. Future research could incorporate gridded habitat data into the same modeling framework to improve estimates of density. This would have the added benefit of being able to generate a total abundance estimate for each species within the Kaʻū Rainforest (Sillett et al., 2012). Moreover, our interaction between year and the quadratic effect of elevation assumes a very particular and smooth relationship with density. Future work could consider more flexible approaches, such as Gaussian processes or additive models, to better capture complex spatiotemporal responses.

### Management implications

Our findings indicate that different species are limited by different stressors operating across the elevational gradient. Endangered honeycreepers continue to contract into increasingly restricted, high-elevation disease refugia, whereas declines in more malaria-tolerant species suggest that habitat degradation and invasive predators are eroding the ecological integrity of lower-elevation forests. Consequently, the conservation value of lower elevations cannot be judged solely by their suitability for the most malaria-sensitive species. Maintaining native bird communities across the Kaʻū Rainforest may require conserving habitat quality throughout the landscape while simultaneously preserving high-elevation refugia.

Area-based management may be necessary to sustain Hawaiʻi’s endemic forest bird communities under ongoing climate change, expanding avian malaria, and biological invasions. Pronounced upslope contractions and local losses of malaria-susceptible honeycreepers, coupled with declines of malaria-tolerant species at lower elevations, demonstrate that passive protection within existing reserves may be insufficient. Instead, effective conservation might require coordinated management of the ecological processes that now drive extinction risk, including mosquito-borne disease, habitat degradation, and invasive predators, implemented across entire protected landscapes rather than at isolated sites. Mosquito suppression within mid-elevation forests, where malaria risk is increasing but susceptible honeycreepers persist, could help slow or reverse the loss of disease refugia. Emerging tools such as the Incompatible Insect Technique (Zheng et al. 2019; Beebe et al. 2021) and larvicides containing *Bacillus thuringiensis israelensis* (Després et al. 2011; Tetreau et al. 2012) provide promising approaches for reducing *Culex* populations and avian malaria transmission. Continued restoration and expansion of native forest through ungulate exclusion, invasive plant control, reforestation, and efforts to limit the spread and ecological impacts of Rapid ʻŌhiʻa Death may further increase carrying capacity while maintaining continuous elevational habitat for species already restricted to narrow bands. Likewise, broad-scale rat suppression may be an effective, core component of area-based conservation rather than a localized management action (e.g., offshore islets; Hess and Jacobi 2011). Carefully planned rat control, including aerial application of rodenticides where appropriate and socially acceptable, may reduce predation pressure on endemic birds and complement mosquito suppression and habitat restoration. Because these threats interact across the landscape, the greatest conservation benefits are likely to be achieved when these strategies are implemented together rather than independently.

Taken together, these interventions suggest a practical framework for prioritizing avian conservation across Hawaiʻi’s protected forests. Disease mitigation and habitat restoration could be concentrated where malaria-sensitive honeycreepers continue to persist, and broad-scale rat control could simultaneously benefit both endangered and more common endemic species. By aligning area-based conservation with the ecological processes limiting individual species, managers can maximize the effectiveness of mosquito suppression, forest restoration, and predator control while maintaining the integrity and resilience of Hawaiʻi’s native forest bird communities.

## Acknowledgments

1. We thank the many dedicated biologists who collected and processed the distance-sampling and vegetation data. This broad-scale assessment of forest birds would not have been possible without collaboration among the following organizations: the National Park Service Pacific Island Network Inventory and Monitoring Network, the State of Hawaiʻi Division of Forestry and Wildlife, Hawaiʻi Volcanoes National Park, the U.S. Geological Survey, the Hawaiʻi Island Natural Area Reserve System Program, the Three Mountain Alliance, Kupu Hawaiʻi, the Maui Forest Bird Recovery Project, the Plant Extinction Prevention Program, The Nature Conservancy, the U.S. Fish and Wildlife Service; the University of Hawaiʻi at Hilo; and the many generous volunteers without formal institutional affiliation.
2. Philip T. Patton’s postdoctoral fellowship was funded by the Cooperative Ecosystems Study Unit grant number P24AC01114-01.
3. All data were collected under the relevant ethical guidelines for distance sampling surveys.
4. We declare no conflicts of interest.
5. **Philip T. Patton:** Conceptualization, Methodology, Software, Formal Analysis, Writing - Original Draft, Visualization **Seth W. Judge**: Conceptualization, Methodology, Investigation, Data Curation, Writing - Original Draft, Project Administration **J. Andrew Royle:** Conceptualization, Methodology, Validation, Writing - Review & Editing, Supervision, Project Administration, Funding Acquisition **T. Scott Sillett:** Conceptualization, Validation, Resources, Writing - Review & Editing, Supervision, Project Administration, Funding Acquisition
6. The data and code to reproduce this analysis can be found on Anonymous GitHub https://anonymous.4open.science/r/pacn-7A7D/README.md and will be archived at Zenodo.

