## Supplementary material for "Modeling two decades of elevational change in endemic bird densities in Hawaiʻi’s Kaʻū Rainforest": Table S2

This information product has been peer reviewed and approved for publication by the U.S. Geological Survey. Any use of trade, firm, or product names is for descriptive purposes only and does not imply endorsement by the U.S. Government.

### Modeling two decades of elevational change in endemic bird densities in Hawai‘i’s Ka‘ū Rainforest

Philip T. Patton [1], Seth Judge [2], J. Andrew Royle [3] and T. Scott Sillett [1]

*[1] Smithsonian’s National Zoo and Conservation Biology Institute, Migratory Bird Center, Washington, DC*
*[2] National Park Service, Pacific Island Inventory and Monitoring Network, Hawai‘i National Park, HI*
*[3] U.S. Geological Survey, Eastern Ecological Science Center, Laurel, MD*

#### Corresponding author

Philip T. Patton,, Migratory Bird Center, Smithsonian’s National Zoo and Conservation Biology Institute, MRC 5503, 3001 Connecticut Ave. NW, Washington, DC, 20013, USA

Table S2: Predicted density (birds per ha) at over time
Each predicted density and (SE) is at a different elevation, in m

| | 2002 | 2004 | 2005 | 2008 | 2010 | 2016 | 2018 | 2019 | 2024 |
|---|---|---|---|---|---|---|---|---|---|
| ʻŌmaʻo | | | | | | | | | |
| 858 | 9.3 (1.85) | 8.7 (1.69) | 8.3 (1.61) | 7.4 (1.41) | 6.9 (1.30) | 5.5 (1.05) | 5.1 (0.99) | 4.9 (0.96) | 4.1 (0.84) |
| 1200 | 7.2 (0.76) | 7.1 (0.73) | 7.0 (0.71) | 6.8 (0.67) | 6.6 (0.65) | 6.2 (0.61) | 6.1 (0.60) | 6.0 (0.60) | 5.7 (0.60) |
| 1530 | 4.8 (0.27) | 4.8 (0.25) | 4.8 (0.24) | 4.8 (0.21) | 4.7 (0.20) | 4.7 (0.20) | 4.7 (0.20) | 4.6 (0.21) | 4.6 (0.25) |
| 1870 | 2.8 (0.24) | 2.7 (0.22) | 2.7 (0.21) | 2.6 (0.19) | 2.5 (0.18) | 2.3 (0.16) | 2.3 (0.17) | 2.2 (0.17) | 2.1 (0.19) |
| 2210 | 1.4 (0.27) | 1.3 (0.23) | 1.2 (0.21) | 1.1 (0.17) | 1.0 (0.15) | 0.8 (0.11) | 0.7 (0.11) | 0.7 (0.11) | 0.6 (0.11) |
| ʻApapane | | | | | | | | | |
| 858 | 39.0 (4.46) | 38.0 (4.20) | 37.5 (4.08) | 36.0 (3.78) | 35.0 (3.61) | 32.3 (3.30) | 31.5 (3.25) | 31.0 (3.23) | 29.0 (3.21) |
| 1200 | 32.4 (1.87) | 32.9 (1.84) | 33.1 (1.83) | 33.9 (1.80) | 34.4 (1.80) | 36.1 (1.85) | 36.6 (1.90) | 36.9 (1.92) | 38.3 (2.11) |
| 1530 | 24.4 (0.75) | 25.5 (0.72) | 26.1 (0.71) | 27.9 (0.67) | 29.2 (0.67) | 33.4 (0.74) | 34.9 (0.81) | 35.7 (0.85) | 40.0 (1.16) |
| 1870 | 16.7 (0.67) | 17.8 (0.67) | 18.3 (0.68) | 20.1 (0.70) | 21.4 (0.72) | 25.7 (0.85) | 27.3 (0.93) | 28.2 (0.97) | 32.9 (1.27) |
| 2210 | 10.4 (0.90) | 11.1 (0.90) | 11.5 (0.90) | 12.7 (0.90) | 13.5 (0.91) | 16.4 (1.06) | 17.5 (1.16) | 18.1 (1.22) | 21.3 (1.65) |
| Hawaiʻi ʻAmakihi | | | | | | | | | |
| 858 | 2.4 (0.52) | 3.0 (0.69) | 0.5 (0.11) | 1.6 (0.36) | 0.8 (0.16) | 1.2 (0.25) | 3.1 (0.73) | 2.0 (0.40) | 1.1 (0.23) |
| 1200 | 4.3 (0.57) | 5.2 (0.82) | 0.9 (0.11) | 2.9 (0.37) | 1.3 (0.18) | 2.1 (0.25) | 5.4 (0.89) | 3.5 (0.37) | 2.0 (0.22) |
| 1530 | 8.1 (0.84) | 10.0 (1.34) | 1.8 (0.13) | 5.5 (0.50) | 2.5 (0.26) | 4.0 (0.32) | 10.3 (1.47) | 6.7 (0.36) | 3.8 (0.24) |
| 1870 | 17.0 (2.00) | 20.9 (3.07) | 3.8 (0.30) | 11.5 (1.16) | 5.3 (0.57) | 8.3 (0.75) | 21.6 (3.35) | 14.1 (0.97) | 7.9 (0.62) |
| 2210 | 38.8 (5.88) | 47.7 (8.44) | 8.6 (1.00) | 26.2 (3.52) | 12.1 (1.65) | 18.9 (2.35) | 49.2 (9.08) | 32.0 (3.60) | 18.1 (2.13) |
| ʻIʻiwi | | | | | | | | | |
| 858 | 0.1 (0.03) | 0.0 (0.02) | 0.0 (0.02) | 0.0 (0.01) | 0.0 (0.01) | 0.0 (0.00) | 0.0 (0.00) | 0.0 (0.00) | 0.0 (0.00) |
| 1200 | 0.9 (0.22) | 0.7 (0.18) | 0.6 (0.16) | 0.5 (0.11) | 0.4 (0.09) | 0.2 (0.05) | 0.2 (0.04) | 0.1 (0.04) | 0.1 (0.03) |
| 1530 | 3.7 (0.36) | 3.4 (0.31) | 3.3 (0.29) | 2.9 (0.24) | 2.7 (0.21) | 2.1 (0.18) | 1.9 (0.17) | 1.8 (0.17) | 1.5 (0.17) |
| 1870 | 4.3 (0.65) | 4.2 (0.61) | 4.2 (0.59) | 4.1 (0.55) | 4.0 (0.53) | 3.8 (0.51) | 3.8 (0.52) | 3.7 (0.53) | 3.6 (0.58) |
| 2210 | 1.4 (0.53) | 1.3 (0.48) | 1.3 (0.45) | 1.3 (0.40) | 1.2 (0.38) | 1.2 (0.38) | 1.1 (0.39) | 1.1 (0.40) | 1.1 (0.47) |
| Hawaiʻi ʻĀkepa | | | | | | | | | |
| 858 | 0.1 (0.10) | 0.0 (0.01) | 0.0 (0.00) | 0.0 (0.00) | 0.0 (0.00) | 0.0 (0.00) | 0.0 (0.00) | 0.0 (0.00) | 0.0 (0.00) |
| 1200 | 0.4 (0.23) | 0.1 (0.08) | 0.1 (0.05) | 0.0 (0.01) | 0.0 (0.00) | 0.0 (0.00) | 0.0 (0.00) | 0.0 (0.00) | 0.0 (0.00) |
| 1530 | 0.8 (0.17) | 0.6 (0.11) | 0.5 (0.10) | 0.3 (0.07) | 0.2 (0.06) | 0.1 (0.03) | 0.0 (0.03) | 0.0 (0.02) | 0.0 (0.01) |
| 1870 | 0.9 (0.35) | 0.9 (0.35) | 0.9 (0.35) | 0.9 (0.35) | 0.9 (0.35) | 0.9 (0.36) | 0.9 (0.37) | 0.9 (0.38) | 0.9 (0.40) |
| 2210 | 0.4 (0.41) | 0.4 (0.39) | 0.4 (0.37) | 0.4 (0.34) | 0.4 (0.33) | 0.3 (0.30) | 0.3 (0.29) | 0.3 (0.29) | 0.2 (0.28) |
| Hawaiʻi ʻElepaio | | | | | | | | | |
| 858 | 6.5 (4.64) | 11.5 (8.18) | 14.8 (10.44) | 25.8 (18.43) | 32.3 (23.27) | 31.5 (22.58) | 24.7 (17.51) | 20.9 (14.74) | 5.9 (4.09) |
| 1200 | 2.3 (0.94) | 4.0 (1.60) | 5.1 (2.03) | 9.0 (3.62) | 11.3 (4.63) | 11.0 (4.52) | 8.6 (3.46) | 7.3 (2.89) | 2.1 (0.81) |
| 1530 | 0.5 (0.11) | 0.9 (0.16) | 1.1 (0.19) | 1.9 (0.35) | 2.4 (0.47) | 2.4 (0.48) | 1.9 (0.35) | 1.6 (0.29) | 0.4 (0.10) |
| 1870 | 0.1 (0.02) | 0.1 (0.04) | 0.1 (0.04) | 0.3 (0.08) | 0.3 (0.10) | 0.3 (0.10) | 0.3 (0.08) | 0.2 (0.07) | 0.1 (0.02) |
| 2210 | 0.0 (0.00) | 0.0 (0.01) | 0.0 (0.01) | 0.0 (0.01) | 0.0 (0.02) | 0.0 (0.02) | 0.0 (0.01) | 0.0 (0.01) | 0.0 (0.00) |
| ʻAlawī | | | | | | | | | |
| 858 | 0.0 (0.00) | 0.0 (0.00) | 0.0 (0.00) | 0.0 (0.00) | 0.0 (0.00) | 0.0 (0.00) | 0.0 (0.00) | 0.0 (0.00) | 0.0 (0.00) |
| 1200 | 0.1 (0.04) | 0.0 (0.02) | 0.0 (0.02) | 0.0 (0.01) | 0.0 (0.00) | 0.0 (0.00) | 0.0 (0.00) | 0.0 (0.00) | 0.0 (0.00) |
| 1530 | 0.4 (0.09) | 0.4 (0.08) | 0.4 (0.08) | 0.4 (0.08) | 0.4 (0.08) | 0.4 (0.09) | 0.4 (0.10) | 0.4 (0.10) | 0.4 (0.13) |
| 1870 | 0.7 (0.28) | 0.8 (0.31) | 0.8 (0.33) | 1.0 (0.40) | 1.2 (0.46) | 1.7 (0.69) | 2.0 (0.80) | 2.1 (0.86) | 3.0 (1.26) |
| 2210 | 0.4 (0.40) | 0.3 (0.29) | 0.2 (0.25) | 0.2 (0.16) | 0.1 (0.13) | 0.1 (0.06) | 0.0 (0.05) | 0.0 (0.05) | 0.0 (0.03) |
| ʻAkiapōlāʻau | | | | | | | | | |

| | 2002 | 2004 | 2005 | 2008 | 2010 | 2016 | 2018 | 2019 | 2024 |
|---|---|---|---|---|---|---|---|---|---|
| 858 | 0.0 (0.00) | 0.0 (0.00) | 0.0 (0.00) | 0.0 (0.00) | 0.0 (0.00) | 0.0 (0.00) | 0.0 (0.00) | 0.0 (0.00) | 0.0 (0.00) |
| 1200 | 0.0 (0.00) | 0.0 (0.00) | 0.0 (0.00) | 0.0 (0.00) | 0.0 (0.00) | 0.0 (0.00) | 0.0 (0.00) | 0.0 (0.00) | 0.0 (0.00) |
| 1530 | 0.0 (0.01) | 0.0 (0.01) | 0.0 (0.01) | 0.0 (0.01) | 0.0 (0.01) | 0.0 (0.01) | 0.0 (0.01) | 0.0 (0.01) | 0.0 (0.01) |
| 1870 | 0.0 (0.02) | 0.0 (0.02) | 0.0 (0.02) | 0.0 (0.02) | 0.0 (0.02) | 0.0 (0.02) | 0.0 (0.02) | 0.0 (0.02) | 0.0 (0.02) |
| 2210 | 0.0 (0.00) | 0.0 (0.00) | 0.0 (0.00) | 0.0 (0.00) | 0.0 (0.00) | 0.0 (0.00) | 0.0 (0.00) | 0.0 (0.00) | 0.0 (0.00) |
